## Supplemental figures for "Simulations of particle tracking in the oligociliated mouse node and implications for left-right symmetry breaking mechanics"

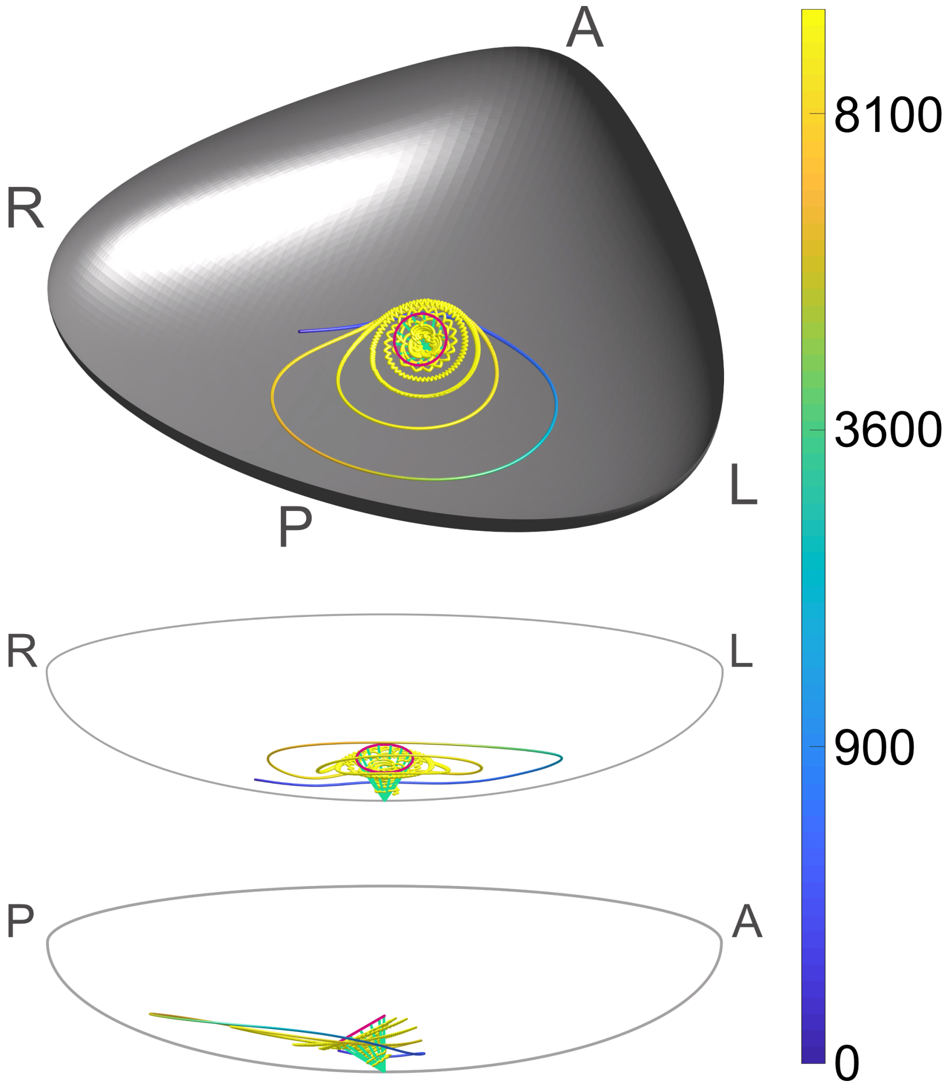


Figure 1: Characteristic particle path for the uniciliated mouse node showing attraction towards the cilia. Time (in cilia beats) is shown according to the colourbar. Side- and front-on views through the node are shown to aid understanding. In each plot the cilium is shown, plotted in green at several points during a beat cycle, with the path traced out by the tip shown in magenta.

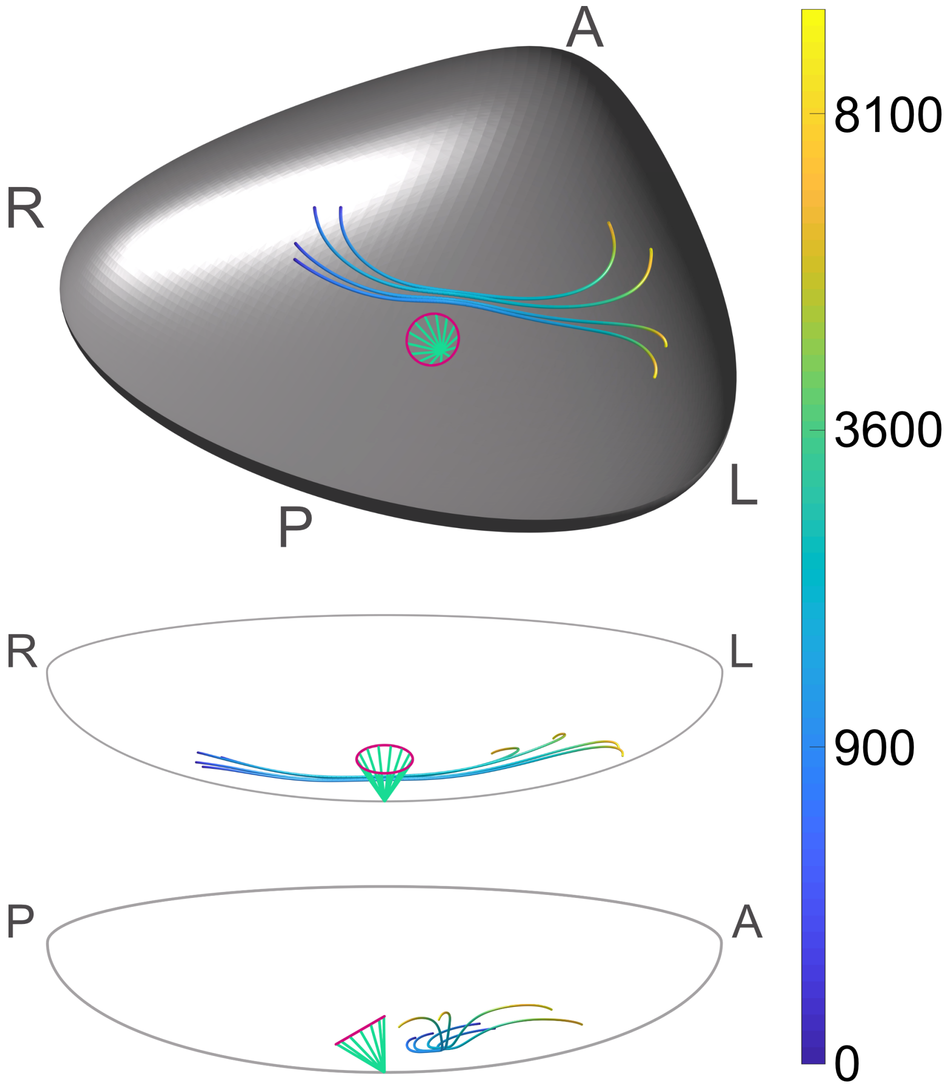


Figure 2: Characteristic particle path for the uniciliated mouse node showing long-range leftwards transport. Time (in cilia beats) is shown according to the colourbar. Side- and front-on views through the node are shown to aid understanding. In each plot the cilium is shown, plotted in green at several points during a beat cycle, with the path traced out by the tip shown in magenta.

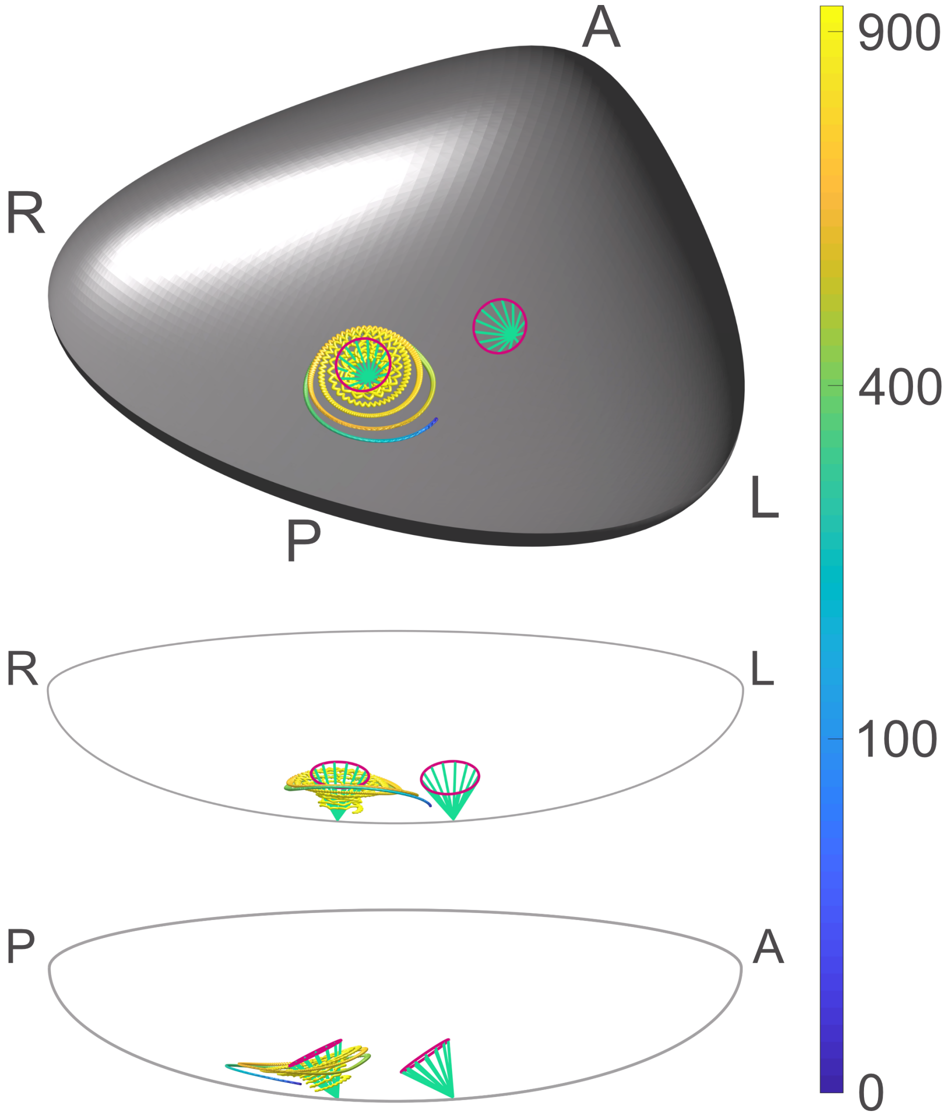


Figure 3: Characteristic particle path for the biciliated mouse node showing attraction towards the rightmost cilium. Time (in cilia beats) is shown according to the colourbar. Side- and front-on views through the node are shown to aid understanding. In each plot the cilia are shown, plotted in green at several points during a beat cycle, with the path traced out by each tip shown in magenta.

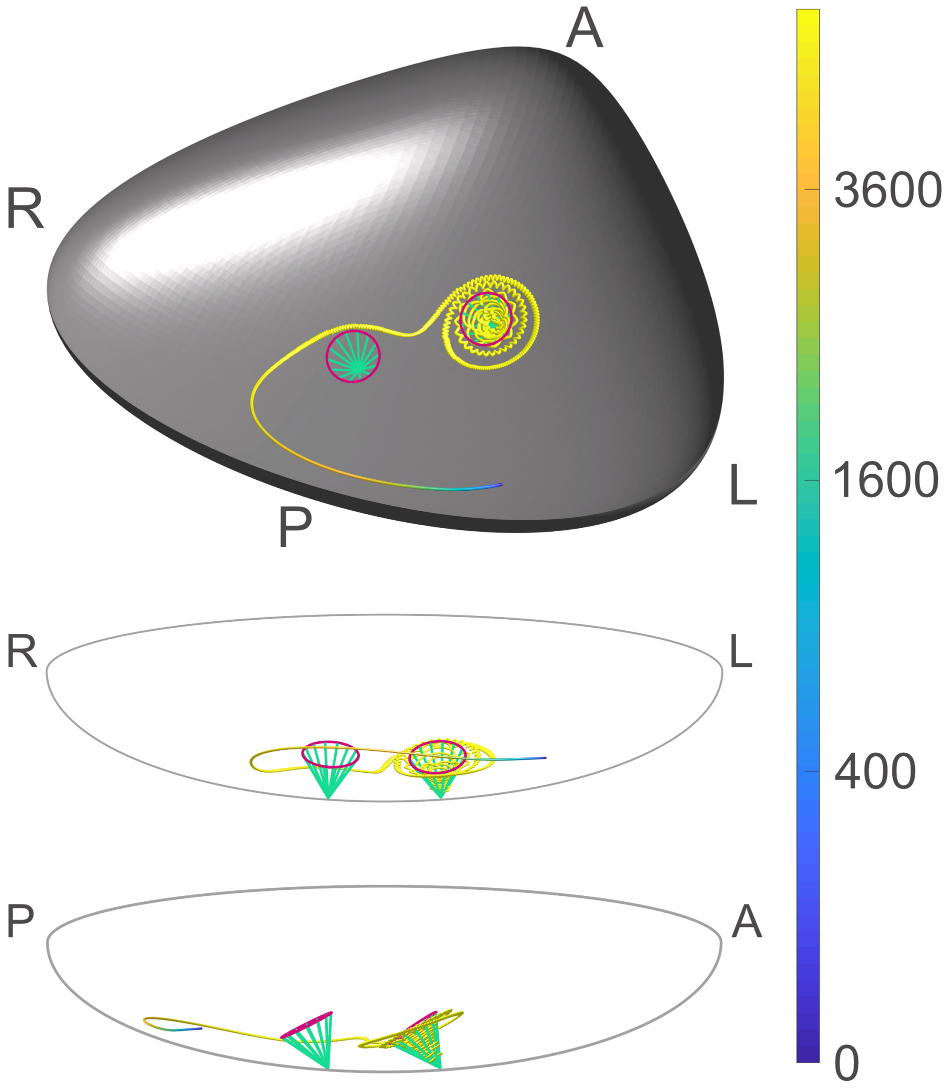


Figure 4: Characteristic particle path for the biciliated mouse node showing attraction towards the leftmost cilium. Time (in cilia beats) is shown according to the colourbar. Side- and front-on views through the node are shown to aid understanding. In each plot the cilia are shown, plotted in green at several points during a beat cycle, with the path traced out by each tip shown in magenta.

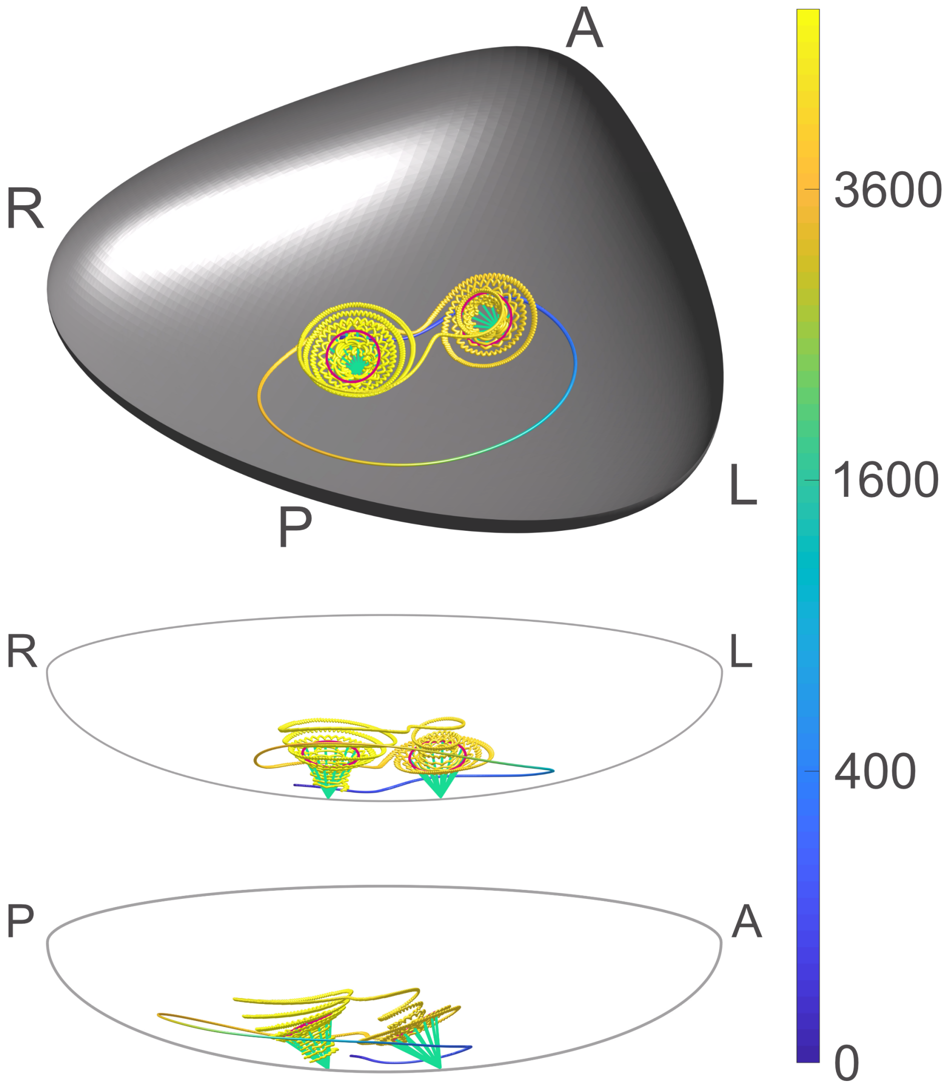


Figure 5: Characteristic particle path for the biciliated mouse node showing particles being passed from one cilium to the other. Time (in cilia beats) is shown according to the colourbar. Side- and front-on views through the node are shown to aid understanding. In each plot the cilia are shown, plotted in green at several points during a beat cycle, with the path traced out by each tip shown in magenta.

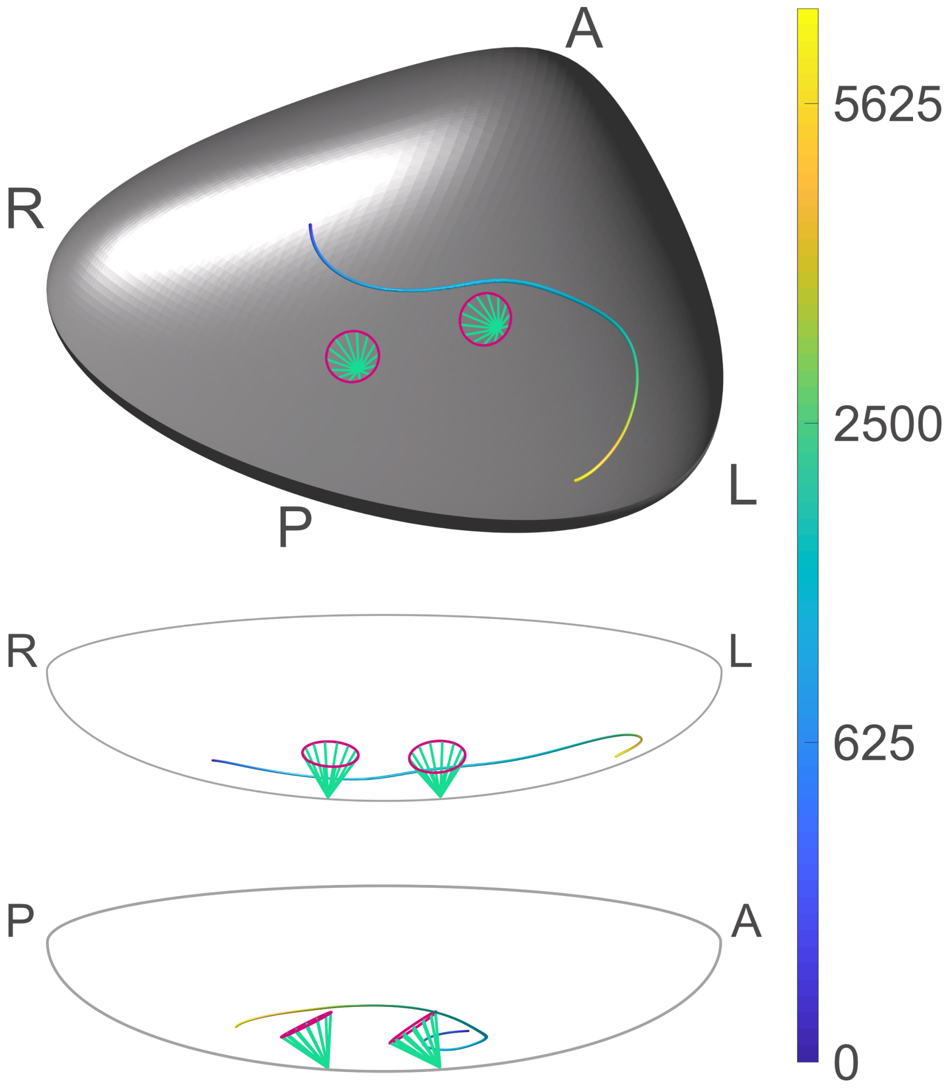


Figure 6: Characteristic particle path for the biciliated mouse node showing long-range leftwards transport. Time (in cilia beats) is shown according to the colourbar. Side- and front-on views through the node are shown to aid understanding. In each plot the cilia are shown, plotted in green at several points during a beat cycle, with the path traced out by each tip shown in magenta.

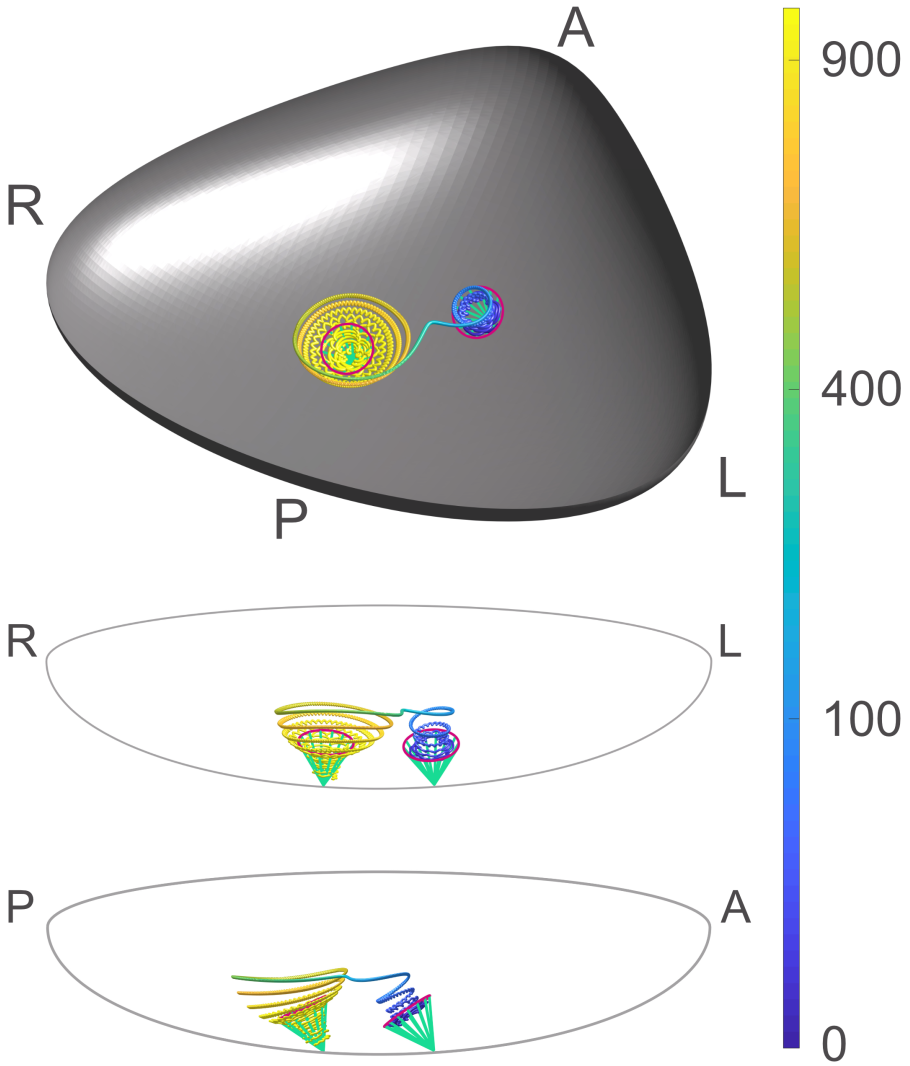


Figure 7: Characteristic particle path for the biciliated mouse node with release directly from the leftmost cilium showing transport from the leftmost to the rightmost cilium. Time (in cilia beats) is shown according to the colourbar. Side- and front-on views through the node are shown to aid understanding. In each plot the cilia are shown, plotted in green at several points during a beat cycle, with the path traced out by each tip shown in magenta.

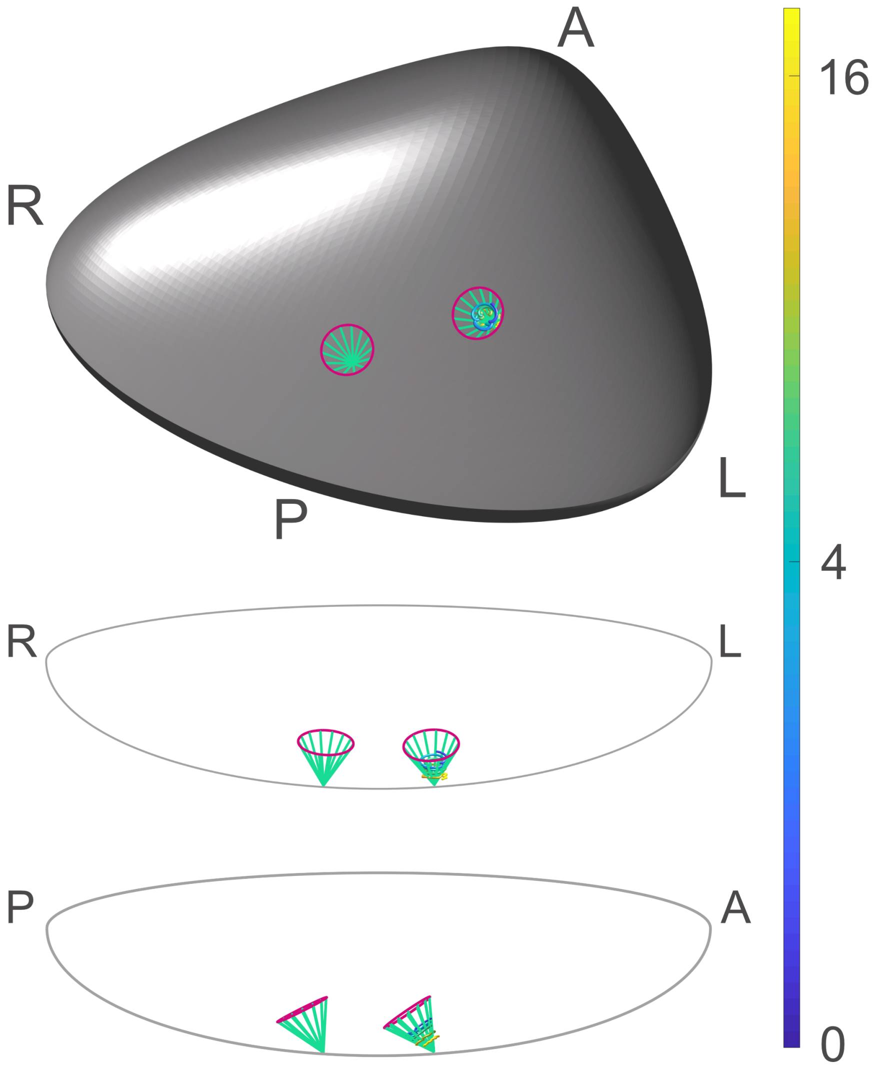


Figure 8: Characteristic particle path for the biciliated mouse node with release directly from the leftmost cilium showing local deposition. Time (in cilia beats) is shown according to the colourbar. Side- and front-on views through the node are shown to aid understanding. In each plot the cilia are shown, plotted in green at several points during a beat cycle, with the path traced out by each tip shown in magenta.

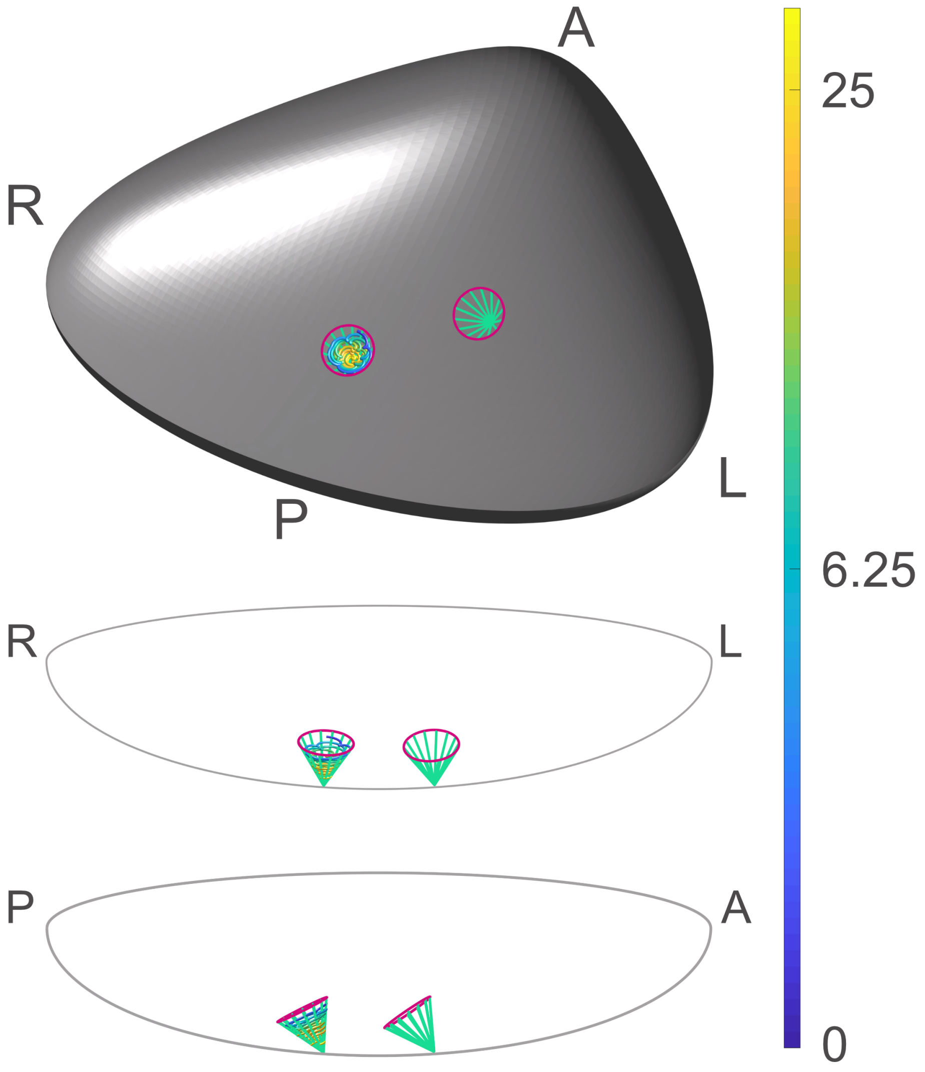


Figure 9: Characteristic particle path for the biciliated mouse node with release directly from the rightmost cilium showing local deposition. Time (in cilia beats) is shown according to the colourbar. Side- and front-on views through the node are shown to aid understanding. In each plot the cilia are shown, plotted in green at several points during a beat cycle, with the path traced out by each tip shown in magenta.

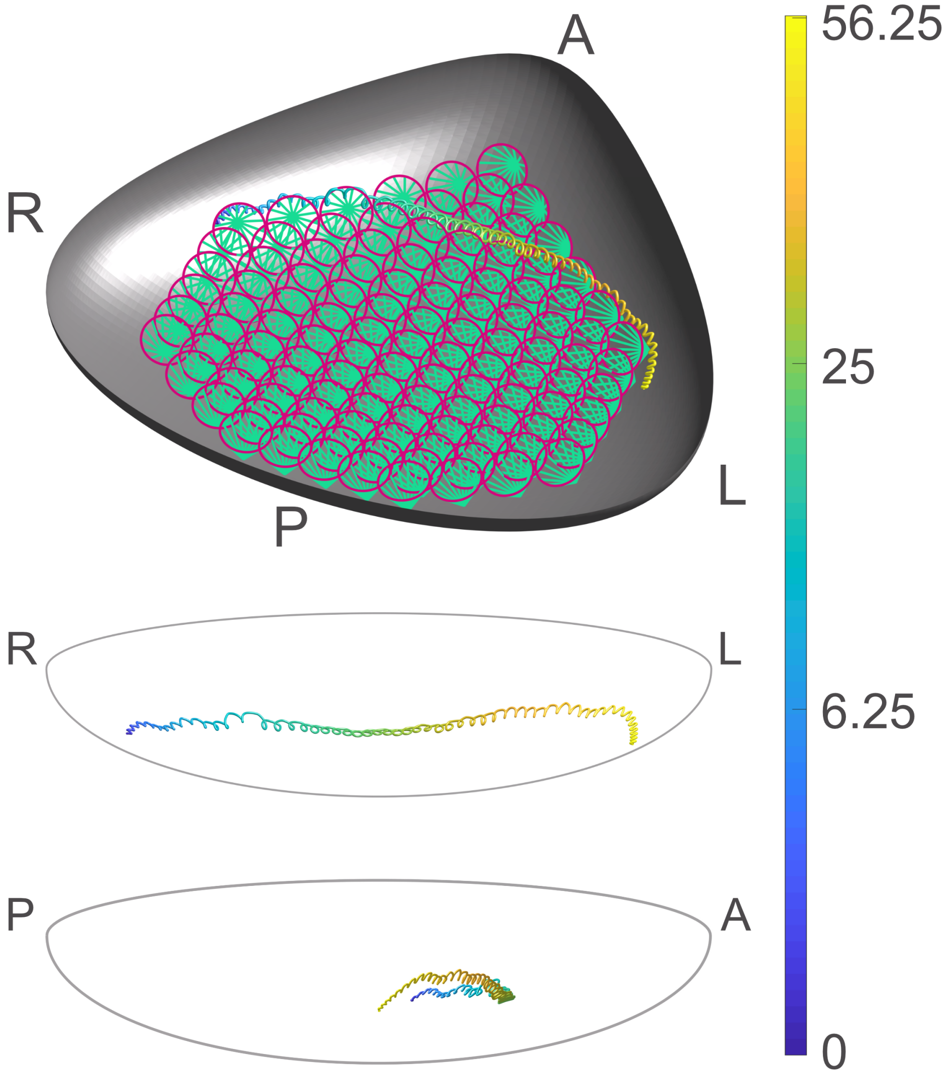


Figure 10: Characteristic particle path for the euciliated mouse node showing long-range leftwards transport. Time (in cilia beats) is shown according to the colourbar. Side- and front-on views through the node are shown to aid understanding. In the tip plot the cilia are shown, plotted in green at several points during a beat cycle, with the path traced out by each tip shown in magenta. The cilia locations are omitted from the side- and front-on views to aid visibility of the particle path.
